# Single-cell DNA cytometry with magnetic- and fluorescence-activated bead sorting

**DOI:** 10.64898/2026.09.12.751168

**Authors:** Seung Won Shin, Sakshi Shah, Chenchen Xia, Atsuki Kawamura, Zhihui Wang, Yuanrong Kang, Nicole Klatt, Iain C. Clark

## Abstract

Many clinically important cell populations are defined by intracellular DNA or RNA signatures, but isolating and profiling these cells remains difficult. Probe-based in situ hybridization strategies that are compatible with cytometry have difficulty resolving low-abundance transcripts, integrated provirus, or single-copy genomic mutations. In contrast, droplet digital PCR achieves single-molecule sensitivity; however, subsequent cell isolation requires specialized microfluidic sorters that are slow and not widely available. Here, we introduce magniFIND-seq, a nucleic acid cytometry platform that combines the sensitivity of digital PCR with commercial magnetic-activated (MACS) and fluorescence-activated (FACS) sorting instruments. Single-cell genomes and transcriptomes are captured in agarose beads, target sequences are detected by digital PCR, and bead-bound amplicons are labeled with magnetic or fluorescent probes. Controlled evaporation shrinks beads from 55 to 20 µm, enabling scalable magnetic separation, fast single-bead FACS sorting, and their combination for high-purity recovery of rare populations. Using magniFIND-seq, we demonstrate multiplexed single-copy detection and FACS-based isolation of simian immunodeficiency virus proviral pol and env targets. Separately, using BCR::ABL1 as a disease-defining target, we enrich chronic myeloid leukemia cells and recover single-cell transcriptomes that resolve tyrosine kinase inhibitor-resistance programs. magniFIND-seq extends the throughput and accessibility of nucleic acid cytometry by engineering compatibility with commercial instruments.

## Introduction

Many disease-relevant cell populations, especially in the omics era, are defined by nucleic acid sequences. For example, cell reservoirs harbor DNA or RNA viruses that evade immune clearance using diverse strategies, including programmed latency [1,2]. In the case of HIV and SIV, the provirus genome persists indefinitely, despite effective suppression of replication, using mechanisms that are largely undefined [3–5]. In cancer, somatic mutations and chromosomal translocations drive malignant phenotypes [6–8] with distinct prognoses and therapeutic responses [9,10]. Isolating and profiling these cells is central to understanding the molecular mechanisms of each disease, yet these populations rarely have unique surface markers suitable for isolation [11,12]. Genome-wide single-cell transcriptome- or open-chromatin sequencing methods are untargeted [13–15] and stochastically capture the specific nucleic acid features (e.g., proviral DNA or fusion transcripts) that define these cells. Consequently, methods are needed to identify and isolate cells according to specific intracellular DNA or RNA sequences.

A solution is to convert nucleic acid targets into sorting markers that enable detection, recovery, and downstream molecular profiling, but current approaches face a trade-off between sensitivity and accessibility. Methods that are compatible with flow cytometry rely on in situ hybridization, often combined with signal amplification, to label intracellular RNA with a fluorescent signal. FISH-Flow [16,17], PrimeFlow [18,19], Probe-Seq [20], and PERFF-seq [21] have shown that target nucleic acid detection can be coupled to cell enrichment and, in some cases, downstream transcriptomic analysis. However, because these approaches depend on fluorescent probes inside fixed or permeabilized cells, their performance is constrained by probe design, intracellular accessibility, and target rarity. In addition, signal amplification is limited, and low-abundance transcripts or single-copy genomic targets can be difficult to detect reliably. In contrast, droplet-based methods such as FADS [22], PACS [23,24], SNAPD [25], and FIND-seq [26–28] use digital PCR to achieve single-copy sensitivity, but they rely on custom microfluidic droplet sorters [29–33] that substantially increase technical complexity and limit throughput and adoption. Thus, a key unmet need is to preserve the single-copy sensitivity of droplet-based PCR while converting its readout into a format compatible with current high-throughput sorting platforms.

We developed magniFIND-seq to combine the sensitivity of PCR-based nucleic acid detection with the accessibility and scalability of commercial MACS and FACS systems. magniFIND-seq retains the core molecular steps of FIND-seq [26], but microfluidic droplet sorting is replaced with magnetic- and/or fluorescent-activated bead sorting. To achieve this, magniFIND-seq combines three technical advances: vortex-based particle-templated emulsification (PTE) for scalable ddPCR [34,35], amplicon tethering within the agarose bead matrix for post-emulsion signal retention, and controlled evaporation-based bead shrinking for compatibility with sorting hardware. Together, these steps convert fluorescent droplets into bead populations that can be processed by standard MACS and FACS platforms. To establish the utility of magniFIND-seq, we demonstrated multiplex detection of pol and env targets from single-copy simian immunodeficiency virus (SIV) proviruses followed by FACS-based isolation. We further applied magniFIND-seq to selectively enrich BCR::ABL1-positive chronic myeloid leukemia cells and recover single-cell transcriptomes that retained biologically meaningful features of tyrosine kinase inhibitor resistance. This establishes magniFIND-seq as a generalizable platform for nucleic acid cytometry (NAC), with potential applications in ultra-rare cell isolation across cancer biology, virology, and immunology.

## Results and Discussion

### Overview of the magniFIND-seq platform

To capture nucleic acids for detection and sequencing, individual cells are co-encapsulated in 55-µm aqueous droplets with lysis buffer and molten agarose conjugated to oligo(dT) primers using a droplet microfluidic device. After lysis, polyadenylated mRNA is hybridized to oligo(dT) primers, and high-molecular-weight genomic DNA remains physically confined within the agarose matrix. The droplets are broken to release hardened agarose beads, and mRNA is reverse-transcribed to generate bead-bound cDNA (**Fig. 1A, Supplementary Fig. 1A**). Three new steps have been introduced in magniFIND-seq to enhance throughput and ensure compatibility with commercial cytometry systems. First, particle-templated emulsification (PTE) on a benchtop vortex mixer replaces microfluidic bead reinjection[34,35], rapidly partitioning millions of beads into PCR droplets within minutes (**Fig. 1B**). This removes a key throughput bottleneck for rare-cell applications and eliminates a complex microfluidic step. Second, PCR amplicons are tethered directly to the agarose polymer, replacing in-droplet soluble TaqMan readout with a stable post-emulsion label (**Fig. 1C**). The bead-bound amplicon allows for biotin-mediated magnetic nanoparticle labeling for MACS and fluorescent labeling for multiplexed FACS sorting. Third, controlled evaporation reduces the agarose bead diameter from 55 to 20 µm (**Fig. 1D**), matching the cell-sized particles that commercial magnetic columns and FACS instruments are designed to efficiently process. Together, these advances convert droplet-based nucleic acid detection into a bead-sorting workflow compatible with high-throughput MACS and FACS instruments that are widely available, extending sequence-defined cytometry beyond specialized microfluidics laboratories.

**Figure 1.**
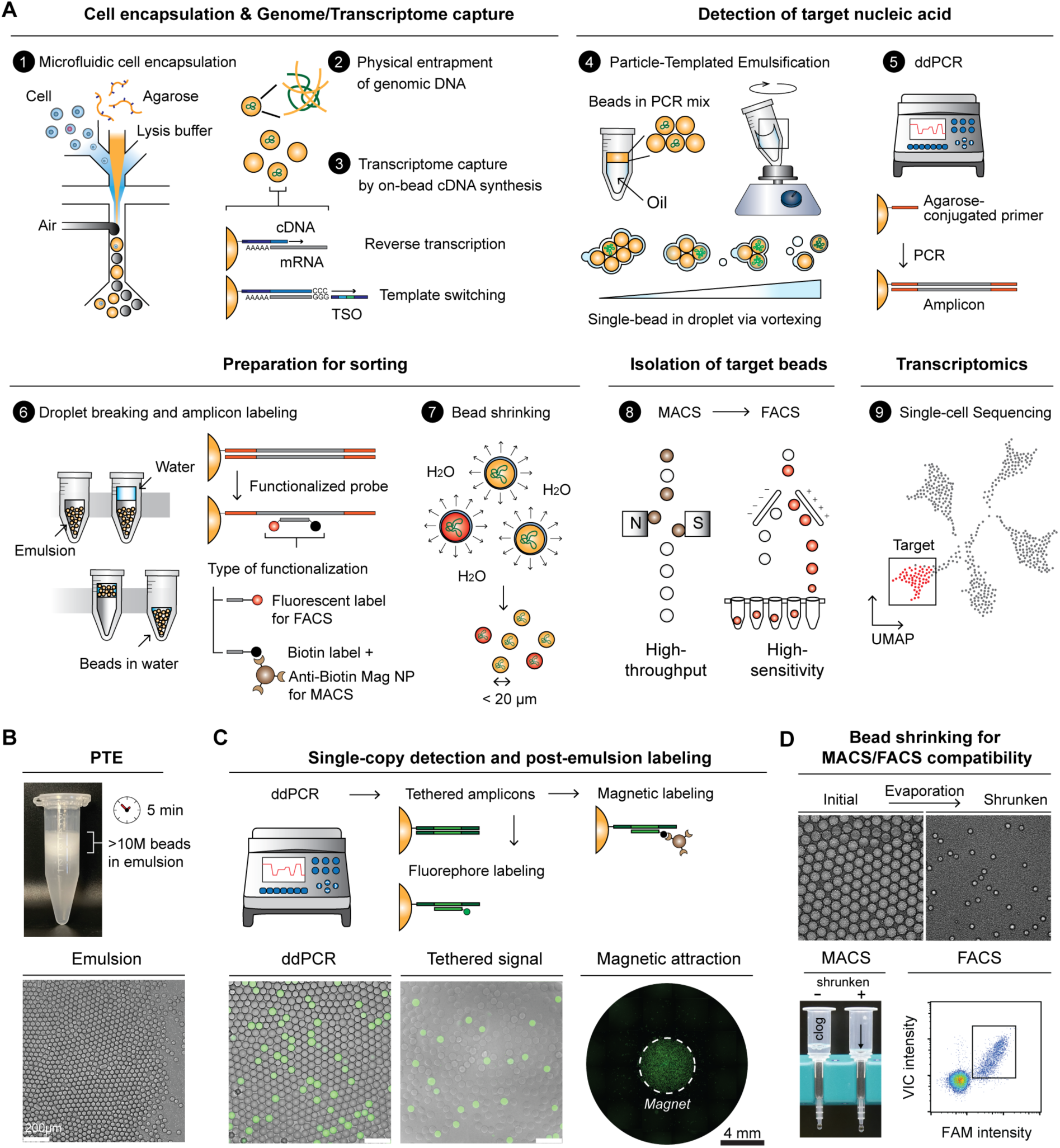
Overview and validation of the magniFIND-seq workflow. (**A**) Schematic of the full workflow. Single cells are encapsulated in agarose droplets containing lysis reagents, resulting in the physical entrapment of genomic DNA and transcriptome capture through on-bead cDNA synthesis (Steps 1-3). Following reverse transcription, the beads are re-emulsified by PTE for ddPCR (Steps 4-5). Target amplicons are tethered to the agarose matrix, labeled after emulsion breaking, and the beads are subsequently shrunk by controlled evaporation (Steps 6-7) before sequential MACS enrichment, FACS isolation, and transcriptomic sequencing (Steps 8-9). (**B**) PTE generates more than 10 million bead-containing droplets within 5 min using a benchtop vortexer. A representative bright-field image shows the resulting bead-templated emulsion. (**C**) ddPCR enables single-copy target detection while retaining amplified products on the agarose beads. After emulsion breaking, bead-tethered amplicons are labeled with fluorophore- or biotin-functionalized probes for fluorescence detection or magnetic capture, respectively. Representative images show bead-associated fluorescence and magnetic attraction following post-ddPCR labeling. (**D**) Controlled evaporation reduces agarose bead diameter from approximately 55 µm to 20 µm. Representative images and size distributions show uniform bead shrinking. After shrinking, the beads are compatible with commercial MACS columns and FACS instruments.

### Amplicon tethering and detection with fluorescent and magnetic labels

TaqMan ddPCR couples amplification to the cleavage of a fluorophore-quencher pair, producing a freely diffusible fluorescent signal. This is not bead-associated and disperses when droplets are broken, precluding recovery of PCR-positive beads by MACS and FACS sorting. To preserve the detection signal through demulsification and allow PCR amplicons to be coupled to both magnetic and fluorescent labels, we tethered PCR amplicons directly to the beads by pre-conjugating the forward primer to agarose. PCR extension generated bead-bound amplicons that were detectable by hybridization with oligonucleotide probes, even after the beads were recovered into the aqueous phase. Tethered double-stranded amplicons were treated with T7 exonuclease to remove the complementary strand and labeled with fluorescent probes for FACS or biotinylated probes for magnetic-column sorting (**Fig. 1C**). We validated this target-detection strategy with three separate PCR assays: mCherry, BCR::ABL1 DNA fusion, and multiplexed detection of SIV pol and env. All three assays successfully generated bead-tethered amplicons, which were detected using amplicon-specific probes (**Supplementary Fig. 1B**).

### Scalable ddPCR with particle-templated emulsification

Preserving single-cell resolution during nucleic acid detection requires compartmentalization of agarose beads into water-in-oil droplets before PCR. Using a bead-reinjection microfluidic device run at 1 kHz (beads/sec) takes approximately 4 hours to encapsulate 1 million cells because each bead is processed serially. In particle-templated emulsification (PTE), a bead suspension is partitioned into droplets in parallel and can generate millions of uniform single-bead emulsions in a single tube in minutes using only a benchtop vortexer. We sought to develop a particle-templated method to eliminate the need for microfluidics and to scale the encapsulation step, but, compared with polyacrylamide beads used in previous PTE studies [34,35], agarose beads required significantly higher vortexing speeds to generate single-bead-in-droplet emulsions. These conditions produced thin aqueous shells that destabilized during thermocycling, causing significant droplet coalescence and loss of single-cell resolution. Recent work has highlighted the unique phase-separation properties of Pluronic F-127 [36–38], a biocompatible triblock copolymer that remains liquid at 4 °C yet forms a rigid, thermo-reversible gel upon warming. We leveraged the properties of F127 to stabilize templated emulsions during thermocycling and maintain detection sensitivity after PTE (**Supplementary Fig. 1C and 1D**).

### Agarose bead size reduction by controlled evaporation

Efficient single-cell lysis, reverse transcription, and droplet PCR occur in droplets larger than 50 µm, whereas commercial magnetic columns and FACS instruments are optimized to process cell-sized particles smaller than 20 µm. Large particles, such as 50 µm agarose beads, caused severe clogging in MACS columns and reduced recovery during FACS. To overcome this limitation, we developed a controlled-evaporation method that uniformly shrinks agarose-containing droplets by approximately 20-fold volumetrically (**Fig. 2**). Agarose beads, re-emulsified by PTE, were heated to melt the agarose matrix within each droplet and subjected to continuous airflow across the air-emulsion interface, which promoted water evaporation from the droplets while maintaining droplet stability (**Fig. 2A**). As water evaporated, the agarose matrix became progressively denser. This process was self-stabilizing: with gentle agitation, denser agarose droplets moved away from the interface, allowing continued evaporation until droplets converged to a uniform final density. Following controlled evaporation, the agarose bead diameter was uniformly reduced to 20 µm (CV 10.4%) (**Fig. 2B-D**), a size that is ideal for MACS- or FACS-based isolation. The shrunken beads also retained the thermo-reversible properties of agarose and could be melted by heating at 65 °C for 1 min (**Fig. 2E**). SEM imaging showed that the agarose structure was maintained, but the gel pore size decreased (**Fig. 2F**). Importantly, oligonucleotide probes remained hybridized throughout the bead-shrinking process (**Fig. 2G)**.

**Figure 2.**
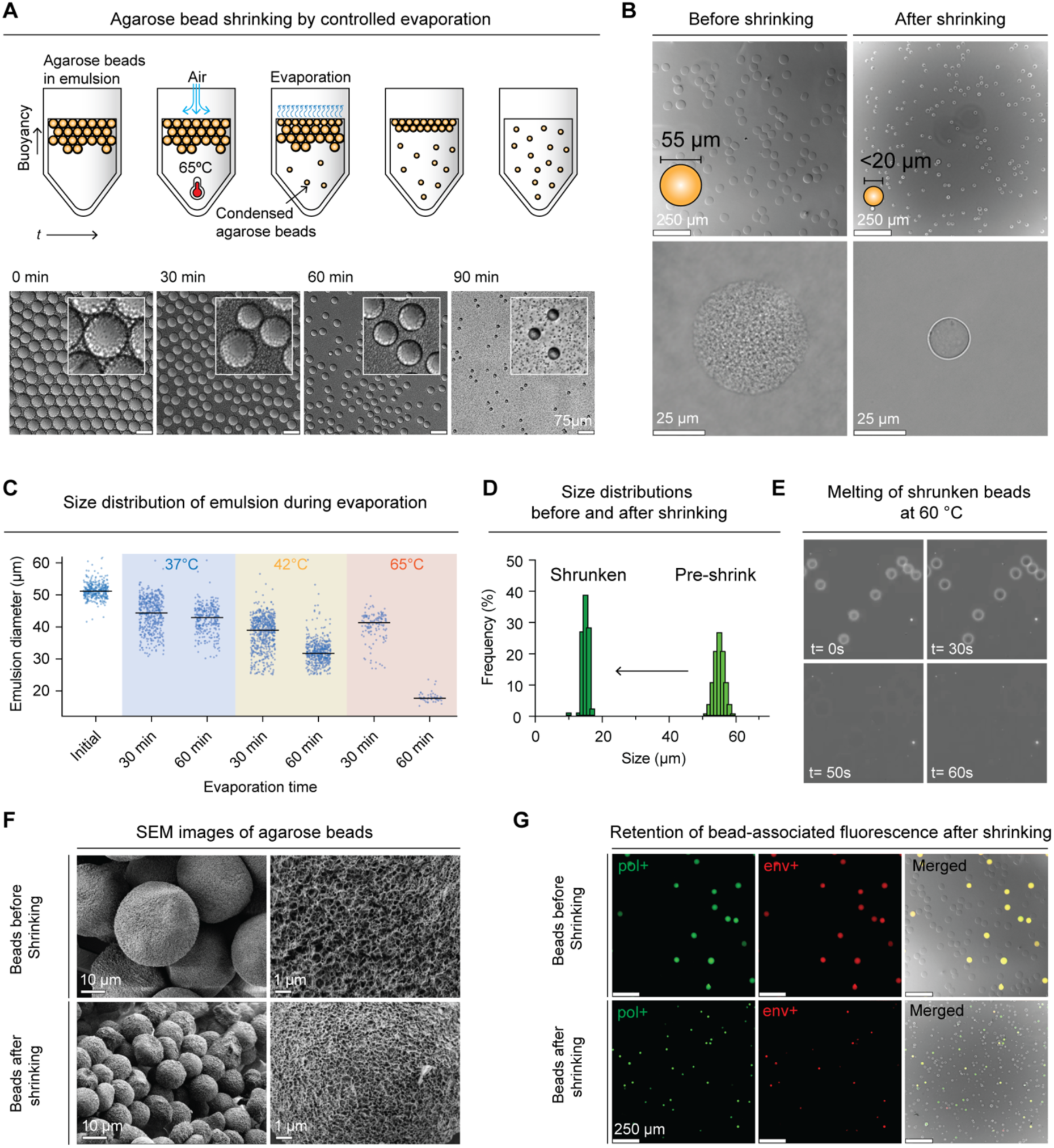
Controlled evaporation enables uniform agarose bead shrinking while preserving agarose bead properties and signal retention. (**A**) Schematic and time-lapse images of agarose bead shrinking by controlled evaporation. Emulsified agarose beads were heated with airflow at 65 °C, promoting water evaporation from droplets at the air–emulsion interface. As evaporation proceeded, agarose beads densified and decreased in size, yielding uniformly shrunken beads. (**B**) Bright-field images of agarose beads before and after shrinking. Bead diameter was reduced from approximately 55 µm to below 20 µm. Lower panels show representative single beads before and after shrinking. (**C**) Distributions of bead-containing emulsion diameters measured during evaporation at 37, 42, and 65 °C at 30 and 60 minutes. Higher temperatures accelerated the reduction in droplet size. Lines indicate the mean. (**D**) Size distributions of beads before and after shrinking, showing a uniform shift from approximately 55 µm to below 20 µm after controlled evaporation. (**E**) Time-lapse images showing the melting of shrunken agarose beads during heating. The beads rapidly lost their structure at 60 °C, indicating agarose maintained its thermal responsiveness after shrinking. (**F**) SEM images of agarose beads before and after shrinking. The porous morphology of the agarose matrix was maintained after shrinking, while the shrunken beads exhibited a denser structure with smaller apparent pores. (**G**) Fluorescence microscopy images showing retention of bead-associated signals after shrinking. Both fluorescent signals remained detectable following controlled evaporation.

### Development of the magnetic-activated bead purification system

MACS is a widely used platform for high-throughput cell enrichment because it is rapid and scalable to large sample volumes. Magnetic cell sorting typically labels target cells with biotinylated antibodies, followed by streptavidin- or anti-biotin-functionalized superparamagnetic nanoparticles. To adapt this scheme to agarose beads, we replaced the antibody with a biotinylated hybridization probe complementary to the tethered PCR amplicon (**Fig. 3A**). Because agarose melts during ddPCR, amplicons are distributed throughout the bead matrix, providing a uniform internal scaffold of labeling sites. Accessing these sites, however, requires magnetic nanoparticles capable of penetrating the agarose pores. We screened several magnetic particles, including Dynabeads MyOne Streptavidin C1 (Thermo Fisher), SuperMag Streptavidin beads (Ocean Nanotech), and Anti-Biotin MicroBeads UltraPure (Miltenyi Biotec), and found that Anti-Biotin MicroBeads UltraPure permeated the bead interior and uniformly labeled tethered amplicons (**Supplementary Fig. 2A**). A dark hue was observed by brightfield microscopy after staining with Anti-Biotin MicroBeads (**Fig. 3A**) and TEM imaging further confirmed the infiltration and retention of magnetic nanoparticles in shrunken agarose beads (**Supplementary Fig. 2B**).

**Figure 3.**
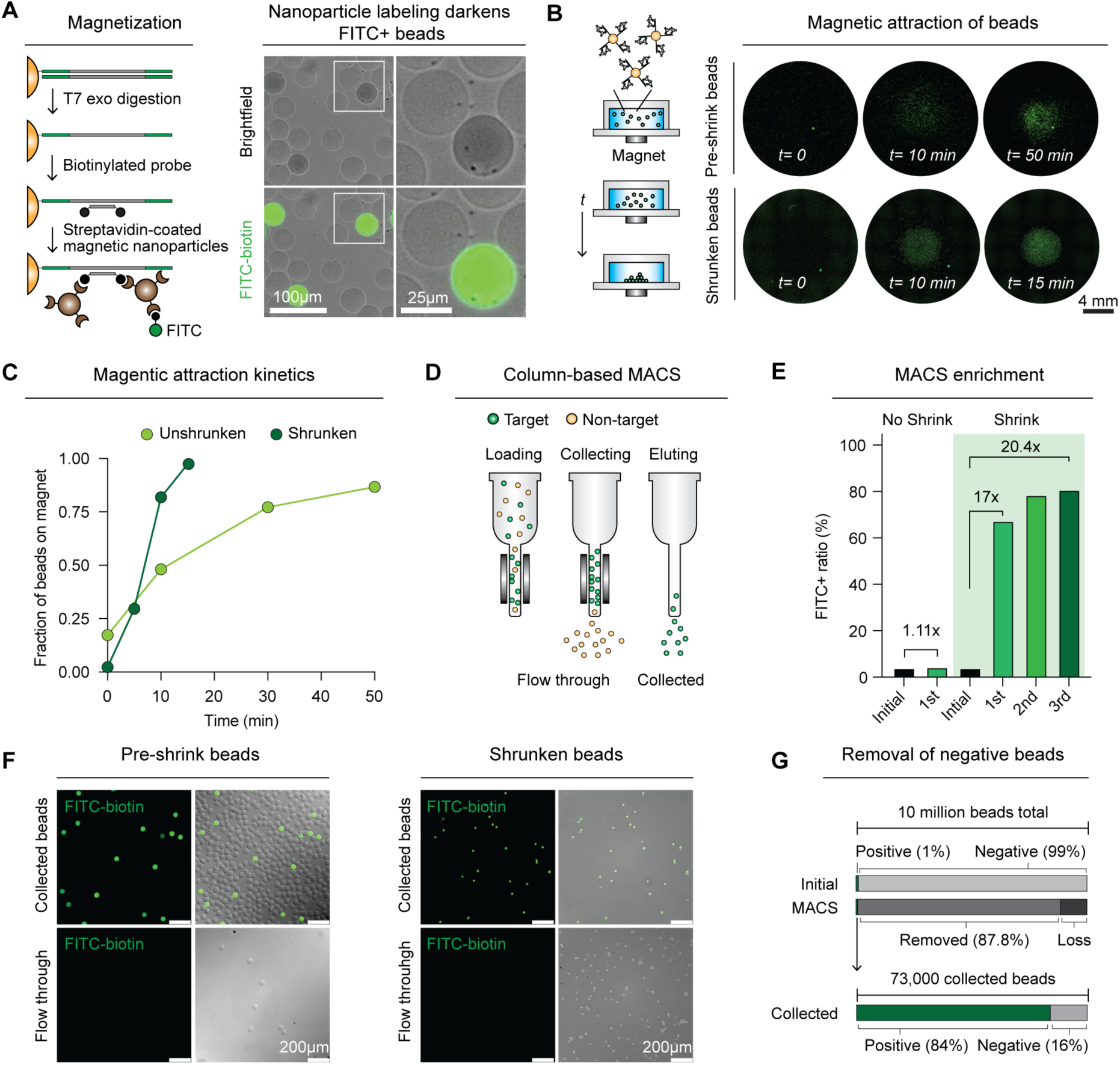
Magnetic nanoparticle labeling and MACS-based enrichment of target agarose beads. (**A**) Schematic of magnetic labeling of bead-tethered amplicons and microscopy images. After T7 exonuclease digestion, single-stranded bead-tethered amplicons were hybridized with biotinylated probes and labeled with streptavidin-coated magnetic nanoparticles. After magnetic nanoparticle labeling, target-positive beads identified by anti-nanoparticle-FITC fluorescence were visibly darker in bright-field images. (**B**) Fluorescence images showing magnetic attraction of labeled beads before and after shrinking. FITC-labeled magnetic beads were placed in wells containing a central neodymium magnet and imaged over time. Shrunken beads accumulated near the magnet more rapidly than beads before shrinking. (**C**) Quantification of magnetic attraction kinetics, shown as the fraction of beads accumulated on the magnet over time. Shrunken beads showed faster magnetic collection than beads before shrinking. (**D**) Schematic of column-based MACS purification. Magnetically labeled target beads are retained during sample loading and washing and subsequently recovered in the collected fraction after removal of the column from the magnetic field. (**E**) Quantification of MACS enrichment over sequential purification rounds. Three rounds of MACS purification increased the FITC-positive target bead population to 80.8%, corresponding to a 20.4-fold enrichment. (**F**) Representative fluorescence and bright-field images of the collected and flow-through fractions obtained using unshrunk and shrunken beads. Compared to 55 µm beads, 20 µm beads passed effectively through the column, and target beads were enriched to high purity. (**G**) Large-scale removal of negative beads by MACS. Ten million shrunken beads containing 1% targets were processed using a commercial MACS column. After a single round of MACS enrichment, the collected fraction contained approximately 73,000 beads, with the target-positive fraction increased to 84%. During the 20-min MACS procedure, 8.78 million negative beads were removed, substantially reducing FACS sorting time.

Having confirmed magnetic labeling of tethered amplicons, we next developed a method to efficiently isolate beads labeled with magnetic nanoparticles. Commercial magnetic separation technologies use a permanent magnet placed beside a tube or well, or a flow-through column containing a ferromagnetic matrix placed in an external permanent magnet. To understand the movement of magnetized agarose beads toward a magnet, we generated PCR-positive beads, labeled the amplicons with complementary biotin-modified oligonucleotides, stained with anti-biotin magnetic nanoparticles, and visualized the accumulation of nanoparticles using FITC-biotin. A 6-mm-diameter neodymium magnet was placed at the center of a 24-well plate to attract the beads, and images were taken over time (**Fig. 3B**). Both 55 µm beads and 20 µm beads accumulated around the magnet, but smaller beads accumulated substantially faster, with more than 90% reaching the neodymium magnet center within 15 min compared to 50 min for larger beads (**Fig. 3C, Supplementary Fig. 3**). Having established that magnetized beads migrate efficiently, we sought to integrate with magnetic separators commonly used for cell isolation. Tube- or well-based magnetic separators are simple and resistant to clogging but typically operate at gradients in the tens to low hundreds of tesla per meter (T/m) and generate their strongest magnetic forces near the magnet-facing vessel wall [39,40]. In contrast, column-based magnetic separators place a packed ferromagnetic matrix within an applied magnetic field. Depending on the matrix geometry, representative local gradients range from approximately 10^3^ to 10^4^ T/m [40]. The matrix also distributes capture surfaces throughout the flow path, reducing particle migration distances from 5-15 millimeters in tube-based systems to the spacing between column matrix elements, typically <100 microns. This much stronger, shorter-range force, together with flow-through washing, makes the column well suited to enriching rare target beads from a large excess of negatives (**Fig. 3D**). The densely packed matrix of column separators, however, accommodates only cell-sized particles, and 55 µm agarose beads quickly clogged commercial Miltenyi columns. In contrast, shrunken beads passed through the column without observable clogging **(Supplementary Fig. 4)**.

To integrate the MACS-column approach, we next evaluated bead enrichment on a commercial MACS column using agarose beads that had undergone ddPCR-based target detection. Agarose beads containing K562 genomes were mixed at ∼4% target abundance, processed by BCR::ABL1 detection ddPCR, and magnetically labeled. Three rounds of MACS purification increased the FITC-positive target bead population to 80.8%, corresponding to a 20.4-fold enrichment (**Fig. 3E and 3F**). To assess large-scale bead processing, we loaded 10 million shrunken beads containing 1% target-positive beads onto the MACS column. After a single round of MACS enrichment, the collected fraction contained ∼73,000 beads, enriching the target population from 1% to 84% while simultaneously removing 8.78 million negative beads in just 20 min (**Fig. 3G**). This substantially decreased the number of beads requiring subsequent FACS sorting. Assuming a FACS sorting rate of 1,000 events/s, MACS pre-enrichment would shorten the estimated FACS sorting time approximately from 2.5 hrs for the initial 10 million beads to 1 min for the collected fraction. Bead shrinking therefore converts agarose beads from column-incompatible to column-ready, transforming millions of beads into a small, target-enriched fraction that can be sorted in minutes. Thus, as in conventional MACS workflows, a single enrichment step removes the vast majority of nontargets, maximizing effective sorting throughput and reducing the sample to a tractable scale so that rare targets can be isolated and sequenced at single-bead resolution.

### Bead sorting using fluorescence-activated cell sorting

MACS provides a simple, high-throughput strategy for target enrichment, but it relies on a single, all-or-nothing magnetic parameter and bulk collection. It cannot resolve multiple markers, discriminate between labeling intensities, or deposit defined numbers of beads. FACS provides a complementary capability by enabling multiparameter analysis, quantitative gating, and precise deposition of defined numbers of beads, including single beads. We therefore sought to adapt commercial FACS instruments for agarose-bead sorting. However, FACS instruments are optimized for cells, and 55-µm agarose beads had several properties that made their isolation by FACS challenging. First, agarose hydrogel beads generated weaker optical scattering (**Fig. 4A, left**), making them hard to distinguish on commercial sorters tested (BD FACSAria Fusion and Bio-Rad S3e), and resulting in low bead recovery after sorting (**Fig. 4B**). Second, their large size destabilized the fluidic stream, increasing dispersion and preventing reliable sorting, particularly of single beads. This was directly observed by measuring the spatial distribution of beads sorted onto glass slides (**Fig. 4C**). Although manual sort-delay adjustment can partially improve large-particle recovery, it does not eliminate the particle-induced instability of droplet breakoff [41,42]. Large nozzle configurations and dedicated large-particle cytometers have been developed for objects outside the practical operating range of conventional cell sorters [43], but their speed and general availability remain limited. Shrinking agarose beads to 20 µm solved both optical and fluidic issues. The denser agarose matrix resulting from shrinkage increased both forward and side scatter, allowing beads to be resolved more clearly as a discrete population by flow cytometry (**Fig. 4A, right**). Smaller beads also passed cleanly through the FACS nozzle, giving more uniform sorts and higher bead recovery (>50% vs ∼10%) (**Fig. 4B and 4C**).

**Figure 4.**
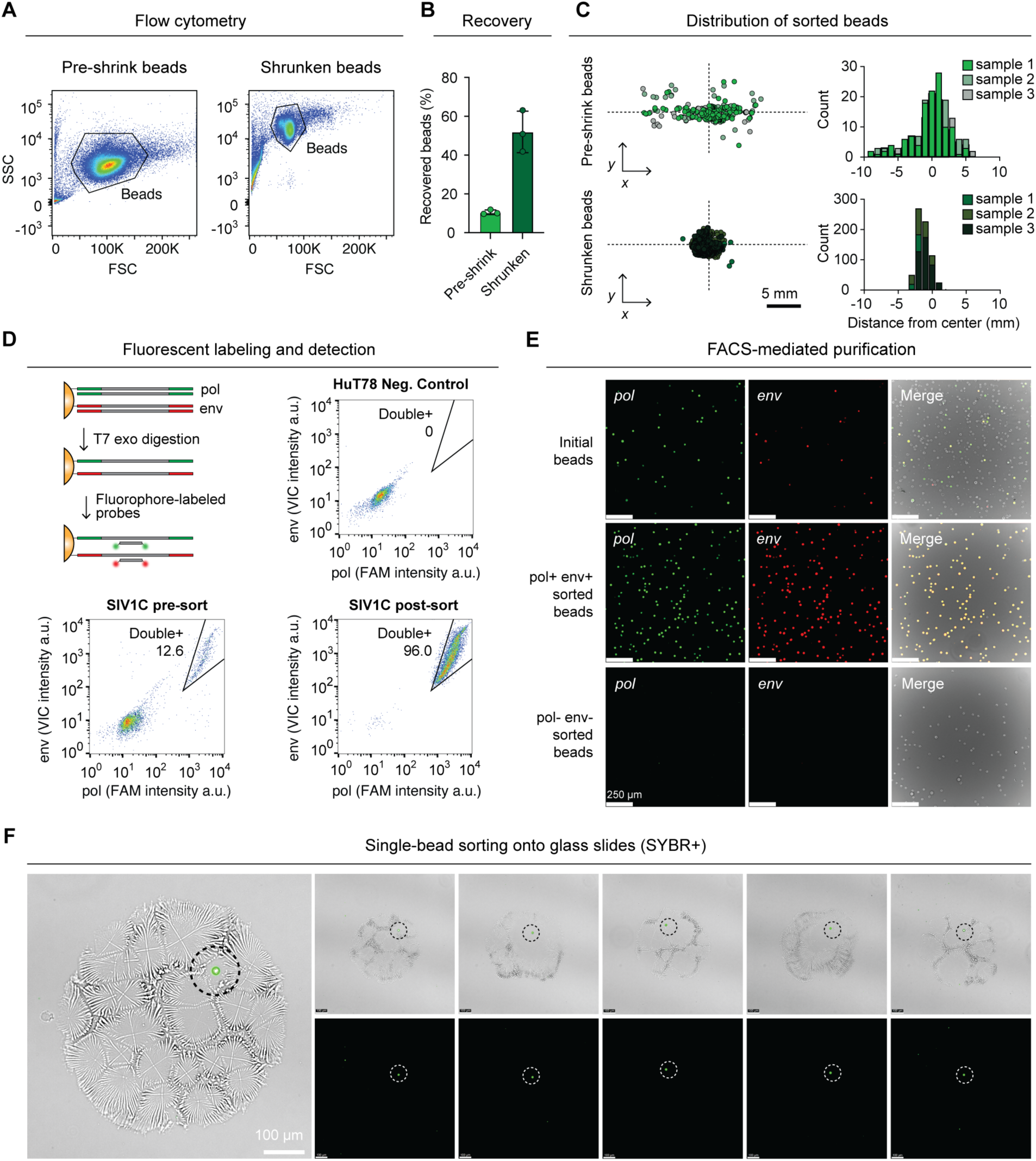
Bead shrinking enables efficient and precise FACS sorting of fluorescently labeled agarose beads. (**A**) Flow-cytometric analysis of agarose beads before and after shrinking. Shrunken beads formed a tight FSC/SSC cluster compared to unshrunken beads. (**B**) Recovery of beads before and after shrinking following FACS sorting onto glass slides. Shrunken beads showed substantially higher recovery than beads before shrinking. Error bars indicate the standard deviation (SD). (**C**) Spatial distribution of beads sorted onto glass slides. Left: shrunken beads were deposited within a smaller area than unshrunken beads, indicating improved sorting precision. Right: distribution of sorted beads in the x-direction from the center. (**D**) FACS gating for multiplexed SIV target detection. SIV1C-derived beads showed a distinct pol+ env+ double-positive population, whereas HuT78-derived negative-control beads showed minimal signal. Post-sort analysis confirmed strong enrichment of the double-positive population. (**E**) Fluorescence microscopy validation of FACS-mediated purification. Sorted pol+ env+ double-positive beads were enriched for both fluorescent signals, whereas sorted double-negative beads showed minimal labeling. (**F**) Single-bead sorting onto glass slides. Representative bright-field and fluorescence images show individually sorted SYBR-positive shrunken beads, confirming single-bead sorting capability.

After demonstrating that agarose beads are compatible with fluorescence-activated sorting, we next validated that magniFIND-seq was capable of multiparameter detection and single-bead sorting precision. These capabilities are especially important for studying heterogeneous cell populations differentiated by more than one nucleic acid sequence. For example, host-integrated human immunodeficiency virus (HIV) and SIV proviruses can contain deletions, hypermutations, and point mutations that render the proviruses incapable of producing infectious virions [44]. Differentiating between defective and intact proviruses and profiling the host cells that harbor these genotypes is an important goal of the field, as it may reveal unique vulnerabilities of cells harboring replication-competent virus. The intact proviral DNA assay (IPDA) is a 2-plex ddPCR method that has become a standard tool for measuring the size of the provirus reservoir [4,45]. To demonstrate sensitive multi-region detection, we used SIV1C cells that harbor a single copy of the SIV provirus, with HuT78 cells as the negative control. Multiplexed amplification using the bead-tethered SIV IPDA assay, followed by labeling with complementary fluorescent probes targeting pol and env sequences and double-positive sorting, enriched pol+ env+ beads to high purity. This was confirmed with post-sort flow cytometry (**Fig. 4D)** and microscopy analysis of the sorted beads (**Fig. 4E**). Cell populations defined by multiple nucleic acid sequences are often heterogeneous and require single-cell sequencing. To confirm that shrinking beads to 20 µm supports single-bead recovery, we sorted fluorescent beads onto glass slides and imaged them, observing reliable single-bead deposition (**Fig. 4F**). Together, these results demonstrate that bead shrinking is critical for agarose bead detection, stream stability, and single-bead recovery on commercial FACS instruments.

### Scalable nucleic acid cytometry using tandem MACS and FACS

After integrating FIND-seq with commercial sorting hardware, we evaluated whether the two enrichment strategies could be used in series to selectively isolate transcriptomes from rare cells carrying a disease-defining nucleic acid marker. Combining scalable, lower-purity MACS enrichment with high-purity single-bead FACS sorting addresses the two competing demands of rare-cell isolation: processing a large excess of non-target cells and resolving the rare targets at single-cell resolution. We focused on BCR::ABL1, an oncogenic fusion gene that defines chronic myeloid leukemia [46,47]. Because BCR::ABL1 arises from a chromosomal translocation and is detected at the DNA or fusion-transcript level, it cannot be directly isolated using conventional surface-marker-based sorting. We designed a ddPCR assay to amplify the genomic translocation junction and modeled the presence of a rare subclone by spiking BCR::ABL1-positive K562 cells into a background of mouse EL4 cells at defined ratios (**Fig. 5A**). Cells were encapsulated into agarose beads to capture genomic DNA and transcriptomes, followed by ddPCR using primers targeting the BCR::ABL1 fusion. The resulting beads were labeled with biotinylated probes and streptavidin-coated magnetic nanoparticles, and processed by sequential MACS and FACS enrichment. A single MACS alone substantially enriched target beads across all spike-in ratios tested. In the 1% K562 spike-in condition, corresponding to 0.1% K562-positive beads, MACS enriched the target bead population by 132-fold (**Fig. 5B**). In cell-mixing experiments containing 1% K562 spiked-in mouse EL4 cells, this large fold-enrichment, coupled with sequential FACS-based sorting of 100 beads into 96-well plates and transcriptome sequencing, increased human reads from ∼1% to ∼90% (**Fig. 5C and 5D)**. Finally, we incorporated a second-strand synthesis step using a random primer containing the smart-seq3 handle to increase yield after whole transcriptome amplification, sorted single beads, and single-cell sequenced material from individual wells. Of the 45 cells that passed quality control, only 3 cells showed a significant mixed-species signature (**Fig. 5E**). Together, these results demonstrate that magniFIND-seq can enrich rare BCR::ABL1-positive populations and recover their transcriptomes with single-cell resolution.

**Figure 5.**
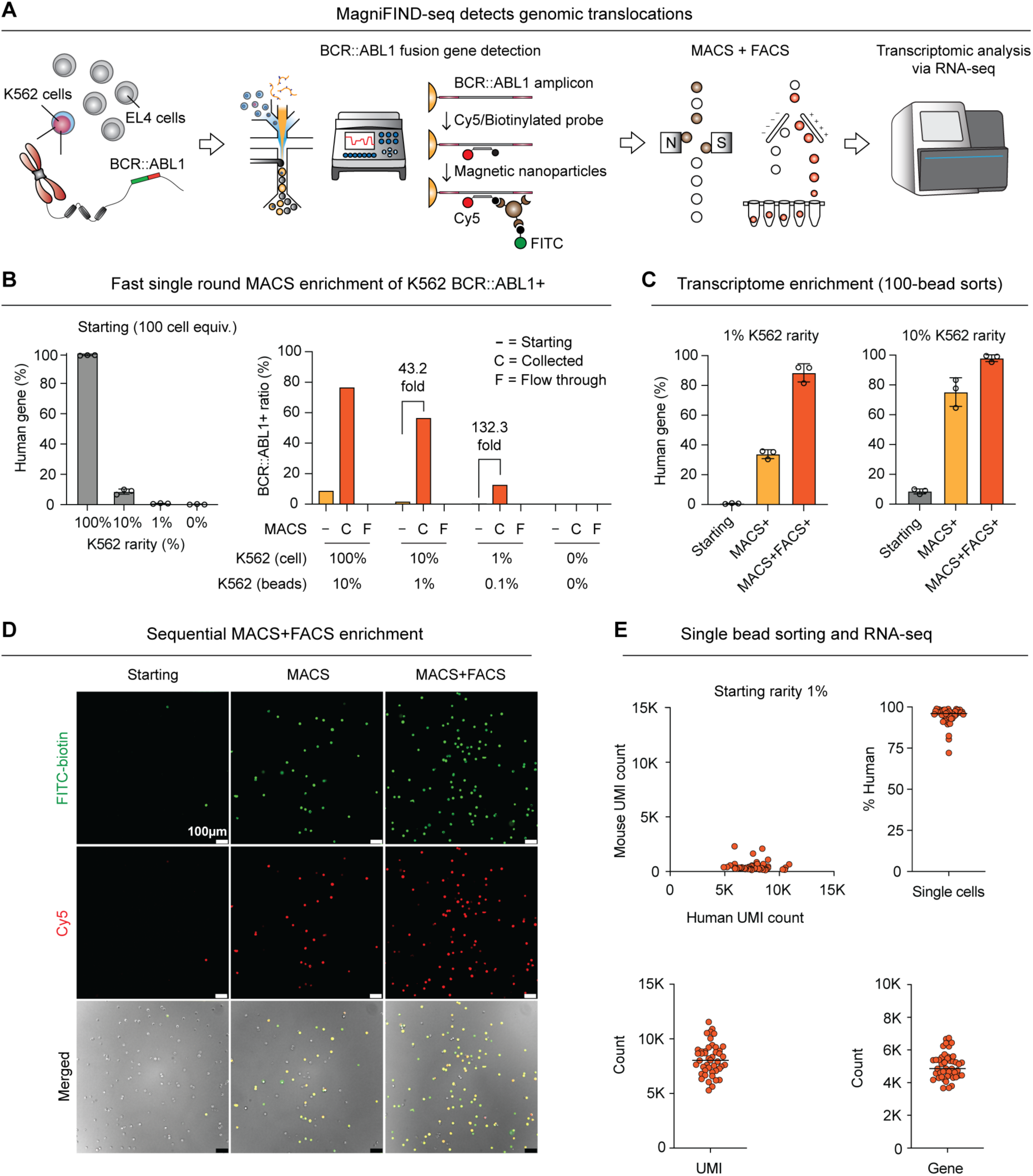
Enrichment of transcriptomes from BCR::ABL1-positive cells by magniFIND-seq. (**A**) Schematic of BCR::ABL1-positive cell enrichment and transcriptome recovery. Human K562 cells harboring the BCR::ABL1 fusion were spiked into a background of mouse EL4 cells and processed by BCR::ABL1-targeted magniFIND-seq. BCR::ABL1-positive beads were labeled with Cy5- and biotin-functionalized probes, magnetically labeled, sequentially enriched by MACS and FACS, and analyzed by RNA sequencing (1000 beads, ∼100 cells). (**B**) Left, initial transcriptomes from control mixtures of 1,000 beads showed human gene fractions corresponding to the K562 input ratios. Right, BCR::ABL1-positive bead fraction quantified in the starting, collected, and flow-through fractions following MACS of different starting rarity levels. Error bars indicate the SD. (**C**) Transcriptome enrichment measured from pools of 100 sorted beads. In samples containing initial K562 cell frequencies of 1% or 10%, MACS increased the fraction of detected human genes, and sequential MACS/FACS enrichment further increased the recovery of K562-derived transcriptomes. Replicate points in (B) and (C) represent three technical replicates generated by sorting 100 beads into separate wells. Error bars indicate the SD. (**D**) Representative fluorescence microscopy images of BCR::ABL1+ beads in the starting population, after MACS enrichment, and after sequential MACS and FACS enrichment. Beads positive for both FITC-biotin and Cy5 signals were progressively enriched after each sorting step. (**E**) Single-bead transcriptome recovery following enrichment of samples containing 1% K562 cells. Human and mouse UMI counts from individually sorted beads. The human gene fraction, total UMI counts, and number of detected genes are shown for individual beads, demonstrating transcriptome recovery at single-bead resolution. Lines indicate the mean.

### Single-cell transcriptomic profiling of TKI-resistant BCR::ABL1-positive cells

Chronic myeloid leukemia is driven by the BCR::ABL1 fusion kinase, the direct molecular target of tyrosine kinase inhibitors (TKIs) [48,49]. However, TKI resistance can arise not only through BCR::ABL1-dependent mechanisms, such as altered BCR::ABL1 activity or expression, but also through BCR::ABL1-independent changes in signaling, survival, stress-response, and drug-adaptation programs [50]. These distinct mechanisms motivate approaches that directly link detection of the disease-defining fusion sequence to single-cell transcriptomic characterization of resistant cells. However, untargeted droplet-based single-cell RNA sequencing, particularly 3′-capture methods, has limited sensitivity for fusion transcripts, especially when the fusion junction lies far from the 3′ end of the transcript. To quantify this limitation, we analyzed a published K562 dataset generated using the 10x Genomics Chromium Single Cell 3′ platform and searched for reads containing the intergenic fusion (see Methods). Out of 5,768 single cells (∼85 million unique reads), only one UMI in one cell identified the BCR::ABL1 fusion (**Supplementary Fig. 5**). We therefore asked whether magniFIND-seq could directly identify cells carrying the disease-defining fusion sequence at the DNA level and recover their transcriptomes for single-cell characterization of TKI resistance.

To model a rare resistant population, we spiked imatinib-resistant K562 cells (K562-r) into human PBMCs at a frequency of 1%. The mixed cells were processed by magniFIND-seq, in which the BCR::ABL1 fusion sequence was detected by ddPCR and enriched by sequential MACS and FACS before being deposited individually into wells for single-cell sequencing (**Fig. 6A**). In total, after sorting, 192 BCR::ABL1-positive single cells were sequenced, of which 182 (94.8%) passed quality-control filtering. The retained cells showed robust transcriptome complexity, with approximately 4,000 UMIs and 2,800 detected genes per cell, while maintaining a low mitochondrial transcript fraction of approximately 4–5% (**Supplementary Fig. 6**). To qualitatively place the recovered K562-r cells in the context of the PBMC background, we jointly visualized the magniFIND-seq-derived transcriptomes with a PBMC single-cell RNA-sequencing reference dataset (**Fig. 6B**). Importantly, in magniFIND-seq, BCR::ABL1-positive cells were identified by ddPCR, their recovery therefore did not depend on capture of the BCR::ABL1 fusion transcript during single-cell RNA sequencing, enabling transcriptomic profiling of sequence-defined cells even when fusion-junction reads were not detected in the downstream libraries (**Fig. 6C and 6D**).

**Figure 6.**
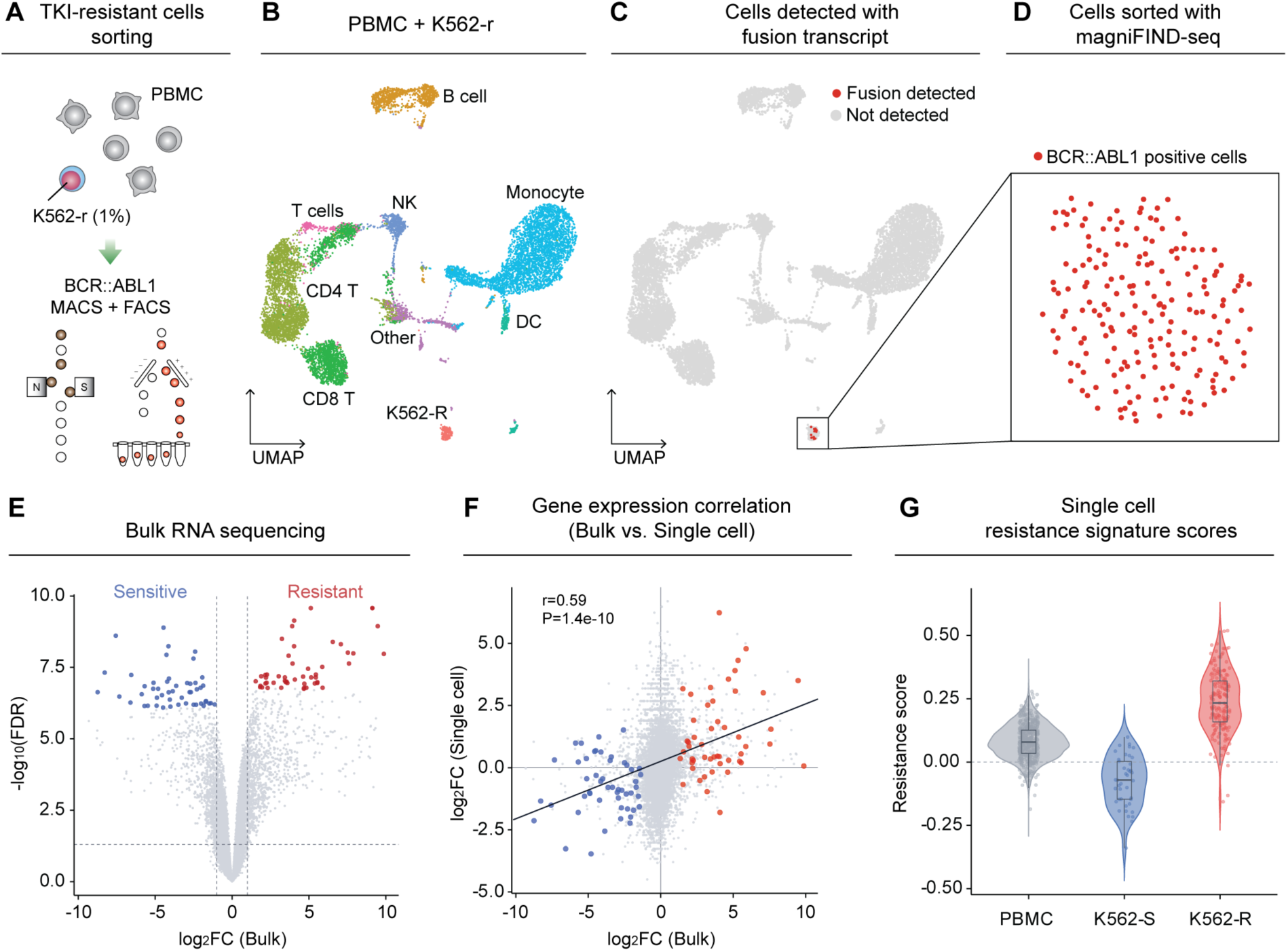
Isolation and single-cell transcriptomic characterization of imatinib-resistant BCR::ABL1-positive cells. (**A**) Schematic of the experimental workflow. Imatinib-resistant K562 cells (K562-r) were spiked into human PBMCs at a frequency of 1% and processed by BCR::ABL1-targeted magniFIND-seq. BCR::ABL1-positive beads were enriched by sequential MACS and FACS and individually deposited into wells for single-cell transcriptomic sequencing. (**B**) UMAP visualization of magniFIND-seq-isolated K562-r cells jointly embedded with a PBMC single-cell RNA-sequencing reference dataset. (**C**) Cells in the integrated UMAP for which sequencing reads supporting the BCR::ABL1 fusion junction were detected. Fusion-supporting reads were identified in 12 of the 182 magniFIND-seq-isolated cells that passed quality-control filtering. (**D**) BCR::ABL1-positive single cells isolated by magniFIND-seq. Each point represents an individually sorted cell selected based on BCR::ABL1 detection independently of fusion-transcript identification by single-cell RNA sequencing. (**E**) Volcano plot of differential gene expression between imatinib-sensitive K562-s and K562-r cells measured by bulk RNA sequencing. The 50 most statistically significant genes enriched in the sensitive and resistant states are highlighted in blue and red, respectively, and used to generate a resistance score. (**F**) Correlation of log_2_ fold changes measured by bulk and single-cell RNA sequencing for genes detected in both datasets. Genes belonging to the sensitive and resistant signatures are highlighted in blue and red, respectively. The solid line indicates the fitted linear relationship (Pearson r = 0.59, P=1.4×10^−10^). (**G**) Resistance scores (see Methods) calculated from the bulk-derived sensitive and resistant gene signatures for PBMCs, K562-s, and K562-r. K562-r exhibited higher resistance scores than K562-s, demonstrating recovery of transcriptomes retaining the imatinib-resistance-associated expression program.

We next assessed whether the recovered single-cell transcriptomes retained biologically meaningful features of TKI resistance. Using bulk RNA-seq, we defined the top 50 most differentially expressed genes between imatinib-sensitive K562 cells (K562-s) maintained under standard culture conditions and K562-r cells, maintained in imatinib-containing medium (**Fig. 6E**). We compared the log_2_ fold changes of these genes between bulk and single-cell datasets and observed a significant positive correlation (r=0.59, P=1.4×10^−10^; **Fig. 6F**). To test this at single-cell resolution, we scored each cell against the bulk-derived resistance signature; magniFIND-seq-sorted K562-r cells scored significantly higher than K562-s cells, confirming that the sorted single-cell transcriptomes retained the bulk-defined resistant state (**Fig. 6G**). Together, these results establish that magniFIND-seq can isolate rare cells defined solely by a disease-associated fusion sequence and recover transcriptomes of sufficient quality to resolve clinically relevant states such as TKI resistance.

## Conclusion

In this study, we developed magniFIND-seq, a bead-based nucleic acid cytometry platform that converts intracellular DNA or RNA targets into magnetic and fluorescent labels compatible with commercial MACS and FACS. By combining vortex-driven particle-templated emulsification, covalent amplicon tethering, and controlled evaporation-based bead shrinking, magniFIND-seq preserves the single-copy sensitivity of droplet PCR while eliminating the need for custom microfluidic droplet sorting. Using this platform, we demonstrated multiplexed detection and sorting of cells harboring SIV provirus and the oncogenic BCR::ABL1 fusion. In the BCR::ABL1 model, magniFIND-seq enriched fusion-positive cells from a mixed human PBMC background by detecting the translocation directly in genomic DNA, rather than the fusion transcript, which is rarely captured in sparse scRNA-seq data. A single MACS step depleted the majority of negative beads, and FACS then recovered rare fusion-positive targets at single-bead resolution, yielding single-cell transcriptomes that retained TKI-resistant signatures. Together, our results establish magniFIND-seq as a generalizable platform for linking single-copy nucleic acid detection to widely available high-throughput cytometry. By providing experimental access to cell populations defined by genomic changes, viral nucleic acids, or other sequence-level features, magniFIND-seq expands the frontier of cell isolation technologies.

## Methods

### Fabrication of microfluidic devices

The bubble-triggered microfluidic device for initial cell encapsulation and lysis in droplets was fabricated using standard photolithography protocols. The bubble-triggered microfluidic device is a trilayer microfluidic device, photomasks for each layer were designed in AutoCAD and printed (Artnet Pro Inc.) [26]. Briefly, a 100-mm silicon wafer (University Wafer # 452) was baked at 150 °C for 10 min before applying SU-8 2025 or 2050 photoresist (Kayaku Advanced Materials) and spin-coating (Laurell Technologies), and soft-baked at 65 °C for 5 min and 85 °C for 15 min on a hot plate. Photomasks for each layer were aligned onto the photoresist-coated silicon wafer using a mask aligner (OAI 206) and exposed to a UV light source of 240 mW/cm^2^. Exposed wafers were developed with SU-8 developer (Kayaku Advanced Materials) for 7 min with gentle agitation, baked overnight at 65 °C, and profiled using a Dektak 3030 stylus profilometer (Bruker). The heights of three layers are: 20 µm, 45 µm, 150 µm, respectively. The patterned silicon wafer was used as a master mold for soft lithography.

Polydimethylsiloxane (PDMS) was mixed with a 10:1 ratio of elastomer and curing agent thoroughly. The mixture was degassed in a vacuum chamber for >30 min, to remove residual air bubbles. The master mold was placed in a clean Petri dish and PDMS was poured onto the center of the wafer. After pouring, the Petri dish was degassed in a vacuum chamber and incubated at 60 °C overnight to cure. Cured PDMS was cut out from the mold and inlet and outlet holes were introduced using a 0.75-mm biopsy punch (World Precision Instruments, Cat. 504529). A microscope glass slide (Fisherbrand, Cat. 12-550C) was cleaned with acetone, isopropyl alcohol (IPA), and distilled water sequentially to prepare for oxygen plasma bonding with the cured PDMS slab. The cleaned glass slide and PDMS slab were placed inside a plasma cleaner (PIE Scientific Tergeo) and treated with oxygen plasma for 30 s at 150 W RF power. After the plasma treatment, the PDMS and glass slide were immediately retrieved and pressed together gently to make tight bonding with further baking at 60 °C overnight. To provide hydrophobicity to the channel, Aquapel (PPG Industries) was introduced to the device channels and incubated for 5 min at room temperature. Residual Aquapel was flushed out with pressurized air and expelled Aquapel was blotted with a lint-free wipe.

### Cell lines

SIV1C, HuT78, K562 and mCherry-CRISPRi K562 lines were cultured in RPMI-1640 (Gibco, Cat. 11875093) supplemented with 10% Fetal Bovine Serum (Avantor, Cat. 89510-186) and 1% Penicillin-Streptomycin (“Pen/Strep” Gibco, Cat. 15140122). EL-4 cell line was cultured in high-glucose DMEM with Glutamax (Gibco, Cat. 10566016) supplemented with 10% FBS and 1% Pen/Strep. K562-r were cultured in RPMI-1640 supplemented with 10% FBS and 1% Pen/Strep, and 1 µM of imatinib (Sigma, Cat. CDS022173-25MG). Cell cultures were maintained at seeding densities and passage ratios according to guidelines from ATCC.

### Agarose conjugation

Agarose was covalently modified with acrydited primers for capture of mRNA and PCR amplicons as previously described [26]. Acrydited primers were ordered dry from Integrated DNA Technologies Inc. (IDT) and conjugated to agarose separately for each capture handle. For conjugation, primers were resuspended to 1 mM in Agarose Conjugation Buffer (0.375 M Tris-HCl Buffer, pH 8.3, Teknova, Cat. T6083). Ammonium persulfate (Sigma, Cat. 215589) was resuspended in Agarose Conjugation Buffer to 10% (w/v) and TEMED (Invitrogen, Cat.15524010) was diluted in Agarose Conjugation Buffer to 10% (v/v). 0.5% SFR allyl agarose (Lucidant Polymers) was resuspended in Agarose Conjugation Buffer in either 15 mL or 50 mL Falcon tubes and heated to 95 °C for at least an hour, until completely molten, with occasional vortexing, and subsequently cooled to 45 °C and incubated for 30 minutes under a vacuum. Acrydited primer, 10% APS, and 10% TEMED were added to the molten allyl agarose to final concentrations of 50 µM, 0.10%, and 0.10%, respectively. The conjugation reaction was incubated under a vacuum for 4 hours at 45 °C, with a repeat addition of 10% TEMED and 10% APS, to a total concentration of 0.20%, and allowed to proceed overnight. The total volume of the conjugation reaction was scaled based on the amount of synthesized oligo received from IDT.

The next day, conjugated allyl agarose was poured into a 10 mL syringe and filtered into a pre-weighed tube with a 0.45-µm syringe filter (VWR, Cat. 76479-020). 1.5% ultra-low gelling temperature agarose powder (Sigma, Cat. A2576) was added to the tube to reach a total agarose concentration of 2% (w/v). The mixture was heated to 95°C for a minimum of 30 minutes, until all agarose was fully molten, and cooled at 4°C for 2 hours. The bottom of the Falcon tube was pierced with a hypodermic needle to release the solidified agarose plug from the tube and solidified agarose was gently transferred to a 500 mL bottle of ultrapure water (Invitrogen, Cat. 10977015) to wash out unconjugated primer. The plug was washed overnight at 4 °C with gentle mixing at 50 rpm on an orbital shaker (New Brunswick Scientific, Innova 2000). The next day, the solidified agarose plug was gently transferred to a fresh bottle of ultrapure water for a second wash. After two washes, the plug was transferred to a Falcon tube. To quantify the concentration of conjugated, modified agarose was heated to 95 °C until molten, and 50 µL agarose was mixed with 3950 µL ultrapure water in triplicate to dilute the modified agarose 80-fold. Mixtures were heated to 95 °C for 15 minutes, cooled to room temperature, and quantified using a Qubit 4 Fluorometer (Invitrogen, Cat. Q33238) and the Qubit ssDNA Assay Kit (Invitrogen, Cat. Q10212). Agarose was conjugated at least three days prior to magniFIND-seq experiments and stored at 4 °C in single-use aliquots in 1.5-mL tube for months.

### Generation of genome- and transcriptome-captured agarose beads

Agarose beads containing an individual cell’s genome and transcriptome were generated using the previously described FIND-seq workflow [26]. In summary, cells were encapsulated into droplets with lysis reagents and oligo(dT)-functionalized molten agarose to capture polyadenylated mRNA and retain genomic DNA within the gel matrix. After gelation, emulsion breaking, and bulk reverse transcription, the recovered beads contained bead-bound cDNA together with physically confined genomic DNA from individual cells. The quality of the bead-bound cDNA was assessed by whole-transcriptome amplification (WTA).

Cells are compartmentalized and lysed inside microfluidic droplets containing Cell Lysis Buffer and molten agarose covalently modified with oligo(dT) primer (IDT, 5’-/5Acryd/TTTTT-TACGAGCATCAGCAGCATACGATTTTTTTTTTTTTTTTTTTTTTTTTTTTTTV-3’). To prepare for cell lysis and mRNA capture, adherent cells were gently detached from plates with 3 mL of 0.25% Trypsin-EDTA (VWR, Cat. E177-100ML) while suspension cells were pipetted from flasks directly into Falcon tubes. Cells were centrifuged and resuspended in 10 mL ice-cold Hanks’ Balanced Salt Solution (Ca-, Mg-, Gibco, Cat. 14175103), filtered with a 40-µm cell strainer (Falcon, Cat. 352340), and counted with a Countess automated cell counter (Invitrogen, Cat. C10228). Meanwhile, a single-use aliquot of oligodT-conjugated agarose was heated at 95°C and occasionally vortexed for a minimum of 30 minutes, until completely molten. Oligo(dT)-conjugated agarose was diluted with unmodified agarose (2% w/v) to reach a primer concentration of 8 µM. 2 mL of 4× Cell Lysis Buffer was prepared fresh by mixing ultrapure water with 20 mM Tris-HCl Buffer, pH 7.5 (Invitrogen, Cat. 15567027), 1000 mM LiCl (Sigma, Cat. L7026-500ML), 1% (w/v) LiDS (Sigma, Cat. L9781-50G), 10 mM EDTA, and 80 U/mL Proteinase K (NEB, Cat. P8107S). Cells were centrifuged and resuspended in ice-cold Cell Resuspension Buffer (Hanks’ Balanced Salt Solution with 18% Optiprep (Sigma, Cat. D1556), and 1% Poloxamer 188, (Gibco, Cat. 24040032) to ensure Poisson-loading of cells into 55 µm droplet compartments with λ = 0.1. Four syringes were prepared as follows: 1-mL syringe with cells in Cell Resuspension Buffer, 3-mL syringe with 4× Cell Lysis Buffer, 3-mL syringe with molten agarose with 8 µM oligo(dT) primer, and 10 mL syringe with 5% (v/v) ionic krytox fluorosurfactant (prepared in-house) in HFE-7500 oil (3M, Novec 7500 Engineered Fluid). Syringes were attached to blunt needles (INSTECH, Cat. LS27) and PE/2 tubing (Scientific commodities, Cat. BB31695-PE/2) and reagents were flowed inside the bubble-triggered microfluidic device using a droplet microfluidic station with a custom syringe heater for the agarose syringe maintained at 90 °C, a custom syringe cooler for the cell syringe maintained at 7-10 °C, a stage heater maintained at 55 °C, and all other reagents kept at room temperature. Flow rates to generate 55-µm droplets were: 600 µL/hour for cells in Cell Resuspension Buffer, 600 µL/hour for 4× Cell Lysis Buffer, 1200 µL/hour for oligo(dT) agarose, 2200 µL/hour for oil, with air pressure maintained at 15-18 psi.

After collecting emulsions, cell lysis was allowed to proceed at room temperature for 2 hours and emulsions were cooled at 4°C for 1 hour. During incubation, buffers for droplet breaking, washing, and reverse transcription were prepared and precooled for at least 1 hour on ice. Buffers were prepared as follows: Droplet Breaking Buffer was prepared by mixing 20% 1H,1H,2H,2H-Perfluoro-1-octanol (PFO, Sigma, Cat. 370533) in HFE-7500 oil, Wash 1 Buffer was prepared by adding 20 mM Tris-HCl, pH 7.5, 500 mM LiCl, 0.1% (w/v) LiDS, and 1 mM EDTA to ultrapure water, Wash 2 Buffer was prepared by adding 20 mM Tris-HCl, pH 7.5 and 500 mM NaCl to ultrapure water, and 5× Reverse Transcription Buffer was prepared by adding 250 mM Tris-HCl, pH 8.3 (Teknova, Cat. T1083), 350 mM KCl (Invitrogen, Cat. AM9640G), and 15 mM MgCl_2_ (Sigma, Cat. M1028) to ultrapure water.

After cell lysis and emulsion cooling incubations, emulsions were immediately broken by removing the oil layer beneath the emulsion, and adding 4 mL of 20% (v/v) PFO and 4 mL of Wash 1 Buffer to the sample tube. Emulsions were incubated at 4°C for 5 minutes with 10 rpm rotation and centrifuged at 100×g for 3 min to separate the oil and aqueous phases. The oil layer was removed and the aqueous phase containing hardened agarose beads remained in the tube. Agarose bead pellets were washed once with Wash 1 Buffer to make up the total volume to 10 mL. After the first wash, any residual breaking oil was removed. Agarose beads were subsequently washed three times with 10 mL of Wash 2 Buffer, twice with 10 mL of 5× RT Buffer, and filtered into a new tube using a 100-µm cell strainer (Fisher Scientific, Cat. 22-363-549). The agarose beads were then transferred to fresh 15-mL tubes containing the reverse transcription mix: 1 mM dNTP mix (ThermoScientific, Cat. R0181), 6 mM MgCl_2_, 1 M betaine (Sigma, Cat. 61962-50G), 7.5% (w/v) PEG-8000 (Promega, Cat. V3011), 2 µM Smart-seq-3-TSO oligo (IDT, 5’-AGAGACAGATTGCGCAATGNNNNNNNNrGrGrG-3’, where rG denotes riboguanine bases), 0.5 U/µL NxGen RNAse Inhibitor (LGC Biosearch Technologies, Cat. 30281), 2 U/µL Maxima H Minus Reverse Transcriptase (ThermoFisher, Cat. EP0751). Sample tubes were incubated at room temperature for 30 minutes with 10 rpm rotation, followed by 42°C for 90 min with 10 rpm rotation. After reverse transcription was completed, 10 mM EDTA was added to each tube and samples were stored overnight at 4°C.

### Whole transcriptome amplification quality check

Agarose beads were washed three times in ice-cold 0.1% Tween-20 in water. A 10 µL aliquot of agarose beads was incubated with 1 µL 100× SYBR Green I dye (Lonza, Cat. BMA50512) for 15 minutes in the dark at room temperature. 10 µL of this mixture was loaded onto a Hemocytometer slide and imaged to quantify the number and ratio of SYBR Green-positive genome-containing beads. A small aliquot of cell-containing agarose beads (“genome beads”) was used to make mixtures of 10 genome beads/µL and 10 µL of the mixture was transferred to a PCR tube strip and mixed with 2× Kapa HiFi HotStart ReadyMix (Roche, Cat. 501965299), and Smart-seq-3-Fwd and Rev PCR primers (IDT, 5’-TCGTCGGCAGCGTCAGATGTGTATAAGAGACAGATTGCGCAA*T*G-3’ and 5’-ACGAGCATCAGCAGCATAC*G*A-3’, 0.5 µM final concentration, * denotes phosphorothioate bonds) for WTA reactions with a total reaction volume of 25 µL. WTA reactions were cycled in a thermal cycler (Eppendorf, Mastercycler X50I) with the following recipe: initial denaturation at 98 °C for 3 min; 14, 16, and 18 cycles of 98 °C for 20 sec, 65 °C for 30 sec, and 72 °C for 4 min; final extension at 72 °C for 5 min, hold at 4 °C. Post-amplification, samples were purified with AMPure XP beads (Beckman Coulter, Cat. A63881) using a 0.8× ratio and quantified using a Qubit 4 Fluorometer with a dsDNA High Sensitivity Kit (ThermoFisher, Cat. Q33230) and TapeStation (Agilent, 4150 TapeStation System) using a D5000 High Sensitivity ScreenTape (Agilent, Cat. 5067-5592) to verify successful cDNA amplification before proceeding with magniFIND-seq detection and enrichment steps.

### Agarose bead re-injection

Agarose beads were prepared in the PCR mixture as described above without F-127. The beads were centrifuged at 1,500 × g for 3 min, and approximately 80% of the upper supernatant was removed. The concentrated bead suspension was thoroughly resuspended and loaded into a syringe connected to the aqueous inlet of a co-flow microfluidic device [26]. QX200 Droplet Generation Oil for EvaGreen was supplied through the oil inlet. The bead suspension and oil were introduced at flow rates of 600 µL/hr and 3,000 µL/hr, respectively, to re-emulsify individual agarose beads in water-in-oil droplets.

### Droplet PCR for target nucleic acid detection and amplicon tethering

PCR strips were thermal cycled with the following recipe: A single initial denaturation step of 1 min at 88 °C, 55 cycles of 88 °C for 30 s and 61 °C for 1 minute, a final anneal/extension step at 61 °C for 1 min, and a hold step at 4 °C. For the emulsion PCR, the ramp rate was lowered to 1.6 °C/s. The annealing/extension temperature was adjusted depending on the genotyping PCR assay used.

### Staining of amplicon with functionalized probes

After thermal cycling, emulsion-containing strips were combined by pipetting and transferred into 15-mL tubes. 2 mL of cold 0.1% (v/v) Tween-20 solution and 2 mL of 20% (v/v) PFO in HFE-7500 were added to the emulsions. The tube was gently inverted 10 times or until all the emulsion clumps were broken up and centrifuged at 100×g for 3 minutes. The oil layer at the bottom of the tube was removed, and 8 mL of 0.1% (v/v) Tween-20 solution was added to the tube. The tube was centrifuged at 2000×g for 3 minutes for the washing, and the washing was repeated three times. The agarose bead pellet was resuspended in a buffer containing 0.1% (v/v) Tween-20, 1% (w/v) sodium dodecyl sulfate (Sigma, Cat. 436143), and 1 mg/mL Proteinase K (NEB, Cat. P8107S) and incubated at room temperature for 1 hour with 10 rpm rotation. Bead pellets were then washed three times with 0.1% (v/v) Tween-20 solution. On the third wash, diluted bead pellets were filtered with a 100-µm cell strainer into a pre-weighed 15-mL tube. After centrifugation, all supernatant was aspirated, leaving the bead pellet behind. And the bead volume was calculated by weighing the tube. The bead pellet was incubated with T7 Exonuclease (NEB, Cat. M0263L) with NEB Buffer 4 (NEB, Cat. B7004S) for 45 minutes at room temperature with 10 rpm rotation. After exonuclease treatment, the complementary strand of the bead-bound amplicons was removed, allowing probes with functionalization to hybridize. The beads were washed three times with 0.1% (v/v) Tween-20 in PBS (PBS-T). The bead pellet was then resuspended in 2 volumes of PBS-T containing 1 µM functionalized hybridization probe. The hybridization mixture was incubated at room temperature for 1 hour with 10 rpm rotation, and an excess amount of probe was removed with three washes with PBS-T.

### Magnetic labeling of biotinylated beads

Post-hybridization with biotin-functionalized probe, the beads were resuspended in PBS with 2 mM EDTA and 0.1% (v/v) Tween-20 solution (Separation Buffer) and incubated with 0.1× of the total volume of Streptavidin MicroBeads (Miltenyi Biotec, Cat. 130-048-101) for 30 minutes at 4°C with 10 rpm rotation to magnetically label beads. Unbound magnetic particles were removed by washing the sample twice with 10 mL Separation Buffer and resuspended in 5 mL Separation Buffer. For microscopy and flow cytometry, magnetized agarose beads can be optionally labeled fluorescently by pelleting the sample, leaving behind 2-fold Separation Buffer, and adding 20 µL of Labeling Check Reagent (Miltenyi Biotec, Cat. 130-124-695).

### Magnetic separation of beads

Magnetized shrunken beads were enriched on a Miltenyi LS column (Miltenyi Biotech, Cat. 130-042-401) using the OctoMACS separator (Miltenyi Biotech, Cat. 130-042-109). Prior to column loading, shrunk beads were resuspended in 10 mL of ice-cold PBS-T, filtered through a 40-µm cell strainer, pelleted, and resuspended in PBS-T. The LS column was primed with 1 mL of PBS-T before adding the beads. The 5 mL sample was applied to the column, followed by two sequential washes with 5 mL PBS-T. The column was removed from the magnet, and the collected beads were eluted into a clean 15 mL tube with 4 × 1 mL of PBS-T, followed by a final 1 mL plunge using the supplied column plunger.

### FACS sorting of beads

Shrunken agarose beads were resuspended in PBS-T after bead recovery and washed before sorting. For fluorescence-activated bead sorting, beads were suspended in at least 3 mL of PBS-T and sorted on a BD FACSAria Fusion cell sorter using a 100-µm nozzle. Bead populations were first identified based on forward- and side-scatter signals, and fluorescence-positive populations were gated according to the labeling strategy used in each experiment. For magnetically labeled samples, FITC labeling was used when applicable to identify target beads. For multiplexed fluorescent detection, target-positive beads were gated using the fluorescence signals from hybridized probes, including Cy5- and FAM-labeled probes. Fluorescence-positive bead populations were sorted directly into 96-well plates, with the number of beads deposited per well specified according to the downstream application. For single-bead transcriptome recovery, individual fluorescence-positive beads were sorted into separate wells of 96-well plates for subsequent whole-transcriptome amplification and library preparation.

### Tagmentation and library preparation

Whole-transcriptome amplification products from sorted beads were purified using 0.8× AMPure XP beads and quantified using a Qubit fluorometer with the dsDNA High Sensitivity Kit. Sequencing libraries were prepared following the Smart-seq3 tagmentation workflow with a modified indexing strategy. Briefly, purified WTA products were tagmented using Tn5 transposase loaded only with the N7 adapter, thereby enabling selective amplification of fragments containing the template-switching oligo-derived sequence at one end. After tagmentation, libraries were amplified using Illumina dual-index primers to introduce sample indices and complete sequencing adapters. Amplified libraries were purified with 0.6× AMPure XP beads, quantified, and analyzed on an Agilent TapeStation to confirm library size distribution and quality. Final libraries were pooled at the desired molar ratio and sequenced on an Illumina NextSeq platform using paired-end 100-bp sequencing.

### mCherry assay

For experiments designed to detect cells containing an integrated mCherry-CRISPRi cassette, primers and probes targeting the mCherry gene were ordered dry from Integrated DNA Technologies and resuspended to 100 µM in ultrapure water. The primer sequences were as follows: mCherry-Fwd, 5′-GTCCTCGAAGTTCATCACGC-3′; and mCherry-Rev, 5′-TTCATGTACGGCTCCAAGGC-3′. For amplicon tethering, the mCherry forward primer was also ordered with a 5′ acrydite modification and a poly(T) spacer for conjugation to agarose: 5′-/5Acryd/TTTTTTGTCCTCGAAGTTCATCACGC-3′. A hybridization probe containing biotin modifications at both the 5′ and 3′ ends was used to label agarose beads containing bead-tethered mCherry PCR amplicons: 5′-/5Biosg/CCCGACTACTTGAAGCTGTCCTTCC/3Bio/-3′.

### BCR::ABL1 assay

For BCR::ABL1 detection, primers targeting the BCR::ABL1 fusion junction were ordered dry from Integrated DNA Technologies and resuspended to 100 µM in ultrapure water. The primer sequences were as follows: BCR::ABL1-Fwd, 5′-GATGACCACGGGACACCTTT-3′; and BCR::ABL1-Rev, 5′-AGGGTATTTCTGTTTGGGTATGGA-3′. For amplicon tethering, the BCR::ABL1 forward primer was also ordered with a 5′ acrydite modification and a poly(T) spacer for conjugation to agarose: 5′-/5Acryd/TTTTTGATGACCACGGGACACCTTT-3′. After droplet PCR and T7 exonuclease treatment, bead-tethered BCR::ABL1 amplicons were labeled by hybridization with a target-specific functionalized probe for downstream magnetic enrichment or fluorescence-activated sorting.

### SIV pol/env multiplexed assay

For multiplex detection of SIV proviral DNA, primers and probes targeting the pol and env regions were ordered from IDT. For the pol assay, the primer sequences were 5′-GCAGGGATAGAGCACACCTTTG-3′ (pol-Fwd) and 5′-CTATGGTTTCTACTGAATTTGCTTGTTC-3′ (pol-Rev), and the TaqMan probe was 5′-/FAM/TTTCAGGTGGTGATTCA/MGBNFQ/-3′. For the env assay, the primer sequences were 5′-CCTCAATAAAGCCTTGTGTAAAATTATC-3′ (env-Fwd) and 5′-GTTATTGTTGATTTTGTCAATCCC-3′ (env-Rev), and the TaqMan probe was 5′-/VIC/TGCATTACTATGAGATGC/MGBNFQ/-3′. For amplicon tethering, the forward primers were synthesized with a 5′ acrydite modification and a poly(T) spacer: pol-Fwd, 5′-/5Acryd/TTTTTGCAGGGATAGAGCACACCTTTG-3′; and env-Fwd, 5′-/5Acryd/TTTTTCCTCAATAAAGCCTTGTGTAAAATTATC-3′. After multiplex droplet PCR and T7 exonuclease treatment, bead-tethered amplicons were fluorescently labeled using target-specific hybridization probes: pol-Hyb, 5′-/FAM/TTTCAGGTGGTGATTCA/FAM/-3′; and env-Hyb, 5′-/MAX/GCATCTCATAGTAATGCA/MAX/-3′. The spectrally distinct probes enabled simultaneous identification and fluorescence-activated sorting of beads positive for the SIV pol and env targets.

### Sequencing read processing, alignment, and gene quantification

Paired-end sequencing reads were demultiplexed according to Illumina dual-index barcodes and converted to FASTQ files. Adapter sequences and low-quality bases were trimmed using Trim Galore. Trimmed reads from human and mouse mixture experiments were aligned with STAR to a combined human and mouse reference genome generated from GRCh38 and mm10. For human-only leukemia mixture experiments, reads were aligned to the GRCh38 reference genome using STAR. Gene-level count matrices were generated from aligned reads using featureCounts with the corresponding gene annotation files.

### Raw read- and UMI-based quality control of single-bead transcriptomes

In the K562 spiked-in mouse EL4 cells experiment, to remove high-complexity samples enriched for putative doublets or mixed-cell barcodes in single-cell analysis, we applied a raw read/UMI-based QC filter. For each sample, we used two sample-level metrics: the number of reads assigned to UMI-tagged features and the total deduplicated UMI count. Empirical cutoffs were defined from the sample-level distributions of these metrics. Specifically, the 65th percentile of UMI-tagged features and the 70th percentile of total deduplicated UMI count were used as thresholds. Samples were retained only if they satisfied both criteria: UMI-tagged features ≤ 478,877 and total deduplicated UMI count ≤ 12,149. Samples exceeding either threshold were classified as raw-QC removals.

### Selection of bulk RNA-seq-derived gene signatures

The top50 gene set was derived from differential expression results from bulk RNA sequencing data processed in R using edgeR package. Multiple-testing correction was performed using the Benjamini–Hochberg procedure, as implemented in stats::p.adjust() with method = "BH". Genes with missing or non-finite gene identifiers or FDR values were excluded using dplyr::filter(), and duplicated gene entries were removed using dplyr::distinct(). When required, the analysis was restricted to genes represented in both the differential expression results and the downstream expression dataset. The remaining genes were ranked in ascending order of FDR using dplyr::arrange(), and the 50 genes with the lowest FDR values were selected using dplyr::slice_head(n = 50). FDR was used as the primary gene-ranking criterion, whereas log2 fold-change was retained to characterize the direction and magnitude of differential expression.

### Concordance analysis between bulk and single-cell RNA sequencing

To assess the concordance between bulk and single-cell RNA-sequencing results, genes from the FDR top50 set were matched between the two datasets using gene symbols. K562-s single-cell transcriptomes were obtained from the human–mouse mixture experiment in Fig. 5, in which K562-s cells were processed using the same magniFIND-seq single-cell sequencing workflow. K562-r single-cell transcriptomes were obtained from PBMC spike-in experiments in Fig. 6. Both K562-r and K562-s scRNA-seq data used the exact same workflow. Single-cell differential expression statistics were obtained using Seurat::FindMarkers(), and only genes with valid log2 fold-change estimates in both datasets were retained using dplyr::inner_join() and dplyr::filter(). Fold-change directions were harmonized so that positive values consistently represented increased expression in the resistant condition relative to the sensitive condition. Gene-level concordance was then evaluated by calculating the Pearson correlation between bulk and single-cell log2 fold-change estimates using stats::cor.test(method = "pearson"). The correlation analysis therefore measured agreement in differential-expression effect sizes rather than correlation between raw expression values. Scatter plots were generated using ggplot2::geom_point() and ggplot2::geom_smooth(method = "lm").

### Calculation of cell-level resistance scores

A cell-level resistance score was calculated from the bulk-derived top50 gene set. Genes were separated into resistance-associated upregulated and downregulated signatures according to the sign of the bulk log2 fold-change. Genes not represented in the single-cell expression matrix were excluded before scoring. Module scores for the two signatures were independently calculated from log-normalized RNA expression using Seurat::AddModuleScore(), which compares the average expression of each signature with that of expression-matched control genes. The final resistance score for each cell was defined as the difference between the upregulated signature score and downregulated signature score. Thus, higher scores indicated a transcriptional state more similar to the bulk resistant phenotype, whereas lower scores indicated a more sensitive-like state. Resistance scores were stored in the Seurat metadata and summarized across experimental groups using dplyr. Score distributions and their spatial patterns were visualized using Seurat::VlnPlot() and Seurat::FeaturePlot(), respectively.

## Author Contributions

I.C.C. conceived the study, supervised the work, and acquired funding. S.W.S. and S.S. developed the methods. S.W.S., S.S., C.X., A.K., Z.W., Y.K., and N.K. performed experiments. S.W.S., S.S., C.X., A.K., and I.C.C. prepared the figures. I.C.C., S.W.S., and S.S. wrote the original draft. All authors reviewed and approved the manuscript.

## Acknowledgements

We thank Misun Kang and Reena Zalpuri of the University of California, Berkeley Electron Microscope Laboratory for advice and assistance with electron microscopy sample preparation and data collection. Electron microscopy performed using the Zeiss XB 550 was supported by NIH S10 Grant S10OD030258-01. This work was supported by QB3 Genomics (UC Berkeley, Berkeley, CA; RRID:SCR_022170) and NIH S10 Instrumentation Grant (S10OD018174). This research used the Savio computational cluster resource provided by the Berkeley Research Computing program at the University of California, Berkeley (supported by the UC Berkeley Chancellor, Vice Chancellor for Research, and Chief Information Officer).

## Funding

ICC and SWS were supported by grants from the NIH (R01DA059551, U54 AI170856, R01AI150396).

## Declaration of interests

ICC and SWS have filed a pending patent application based on the technology described in this manuscript.

## Data availability

The raw and processed bulk and single-cell RNA-sequencing data generated in this study have been deposited in the NCBI Gene Expression Omnibus (GEO) under accession numbers GSE342953 and GSE342951, respectively. A version of the code corresponding to this study will be permanently archived in Zenodo and the associated DOI will be provided upon publication. The publicly available PBMC single-cell RNA-sequencing dataset used for data integration was the “10k Peripheral Blood Mononuclear Cells (PBMCs) from a Healthy Donor, Single Indexed” dataset generated by 10x Genomics and processed using Cell Ranger 4.0.0 (https://www.10xgenomics.com/datasets/10-k-peripheral-blood-mononuclear-cells-pbm-cs-from-a-healthy-donor-single-indexed-3-1-standard-4-0-0). The publicly available K562 single-cell RNA-sequencing dataset used for chimeric transcript analysis was SRR5082088, corresponding to GEO sample GSM2406675.

