## Supplemental Information for "Single-cell DNA cytometry with magnetic- and fluorescence-activated bead sorting"

### Contents

**Figure S1.** Validation of bead-bound cDNA amplification and bead-tethered amplicon detection.

**Figure S2.** Screening of magnetic particles for labeling bead-tethered amplicons in agarose beads.

**Figure S3.** Size reduction accelerates magnetic attraction of magnetically labeled agarose beads.

**Figure S4.** Bead shrinking prevents clogging during MACS column processing.

**Figure S5.** Prevalence of putative intergenic chimeric transcripts in the publicly available K562 single-cell RNA-sequencing dataset.

**Figure S6**. Quality-control metrics of magniFIND-seq-isolated K562-r single-cell transcriptomes.

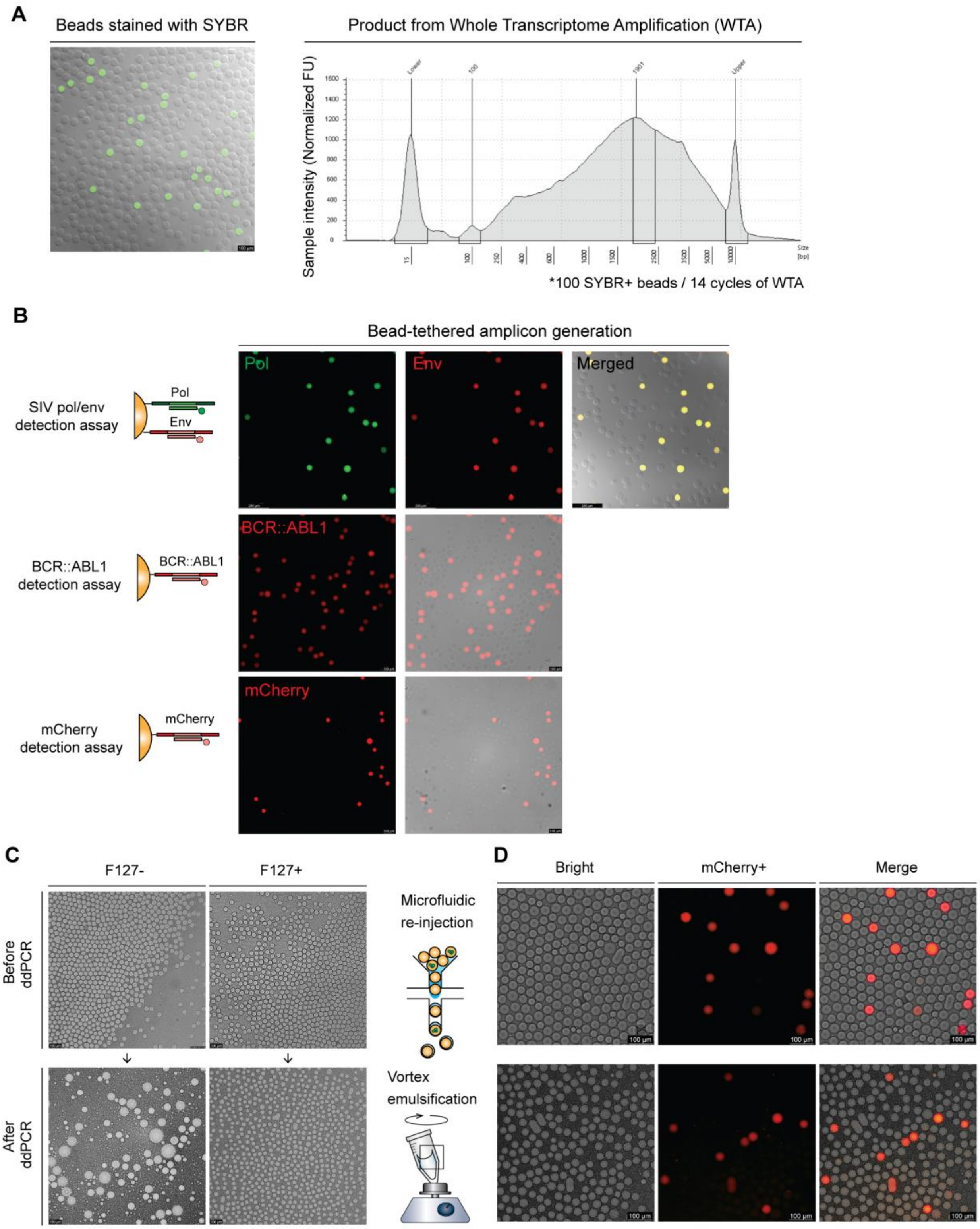
A
Beads stained with SYBR
Product from Whole Transcriptome Amplification (WTA)
Sample intensity (Normalized FU)
*100 SYBR+ beads / 14 cycles of WTA
B
Bead-tethered amplicon generation
SIV pol/env detection assay
Pol
Env
Merged
BCR::ABL1 detection assay
BCR::ABL1
mCherry detection assay
mCherry
C
F127-
F127+
Before ddPCR
After ddPCR
Microfluidic re-injection
Vortex emulsification
D
Bright
mCherry+
Merge
100 µm

**Figure S1. Validation of bead-bound cDNA amplification and bead-tethered amplicon detection.**

(**A**) Validation of genome/transcriptome capture and whole-transcriptome amplification (WTA) from agarose beads. SYBR staining identified genome-containing agarose beads, and WTA products generated from 100 SYBR-positive beads after 14 amplification cycles showed a broad size distribution consistent with successful transcriptome amplification. (**B**) Validation of bead-tethered amplicon generation across multiple target-detection assays. Agarose beads were subjected to ddPCR using agarose-conjugated primers, followed by T7 exonuclease digestion and hybridization with target-specific fluorescent probes. Representative fluorescence microscopy images show multiplexed detection of SIV pol and env targets and single-target detection of BCR::ABL1 and mCherry. (**C**) Stabilization of bead-templated emulsions by Pluronic F-127 during PTE-based ddPCR. Representative bright-field images show bead-containing emulsions generated in the absence or presence of Pluronic F-127 before and after ddPCR thermocycling. In the absence of Pluronic F-127, the emulsion became unstable during thermocycling, whereas inclusion of Pluronic F-127 maintained single-bead occupancy of droplets after ddPCR. (**D**) Comparison of mCherry detection following droplet PCR using microfluidic bead reinjection or vortex-driven particle-templated emulsification. Representative bright-field and fluorescence images show mCherry-positive emulsions generated using both emulsification methods, demonstrating that vortex-driven PTE supports ddPCR-based detection.

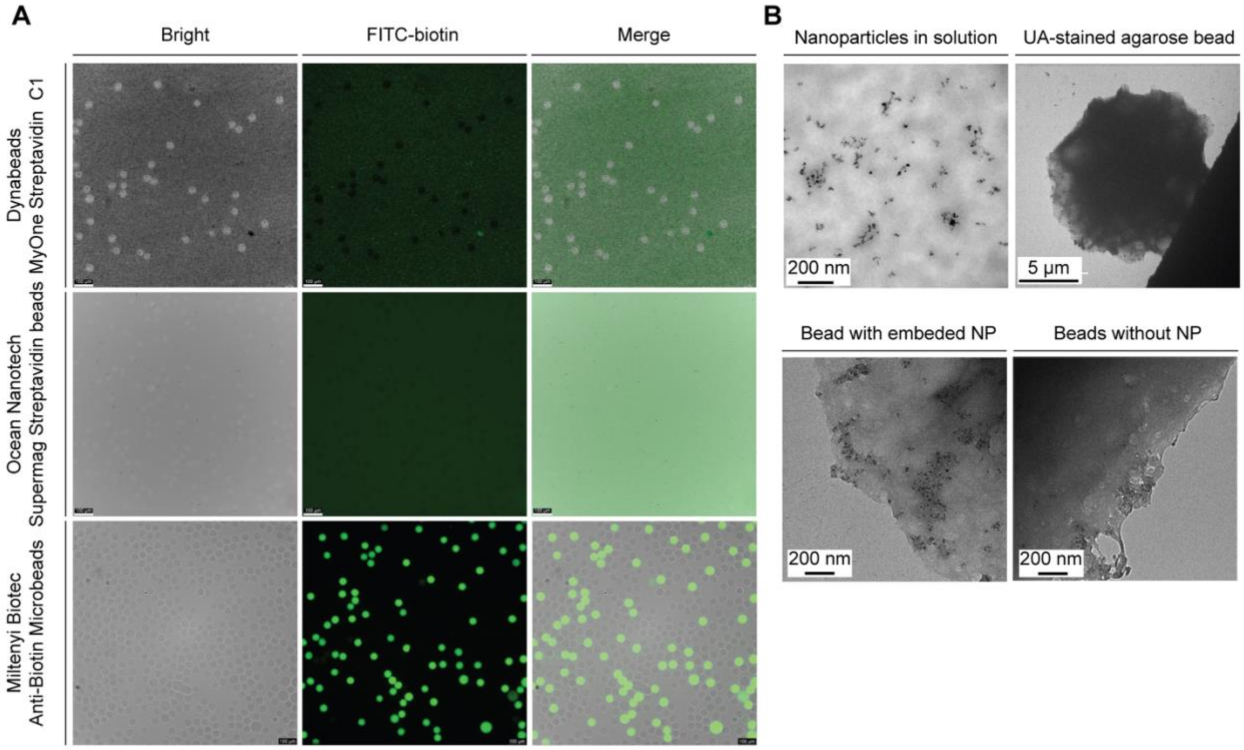


**Figure S2. Screening of magnetic particles for labeling bead-tethered amplicons in agarose beads.**

(**A**) Fluorescence microscopy images of agarose beads labeled with different streptavidin- or anti-biotin-coated magnetic particles. Dynabeads MyOne Streptavidin C1 and Ocean NanoTech SuperMag Streptavidin beads showed limited labeling of agarose beads, whereas Miltenyi Biotec Anti-Biotin MicroBeads UltraPure produced strong and uniform FITC-biotin labeling throughout the bead population. **(B)** Transmission electron microscopy (TEM) images of Miltenyi Biotec Anti-Biotin MicroBeads UltraPure in solution and within uranyl acetate (UA)-stained agarose beads. Magnetic nanoparticles were observed within the agarose matrix after magnetic labeling but were absent from most of the beads, consistent with penetration of the small magnetic particles into the porous agarose matrix.

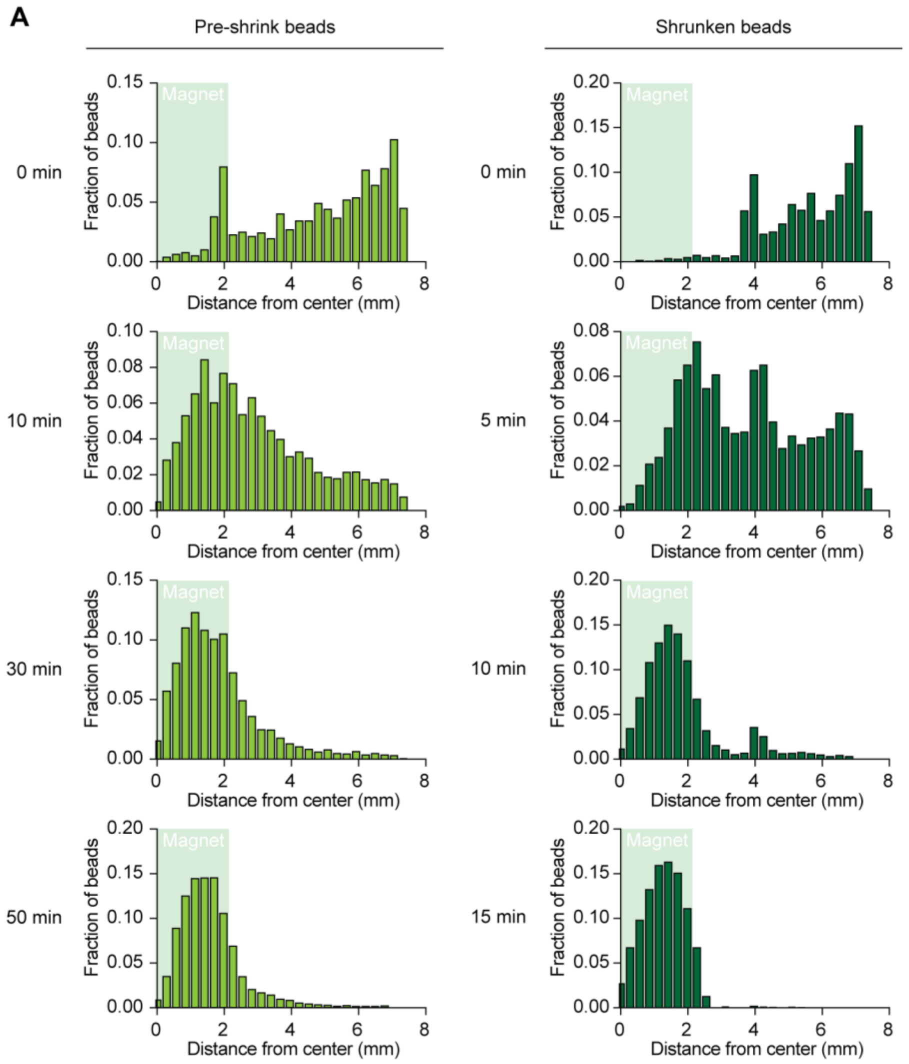


**Figure S3. Size reduction accelerates magnetic attraction of magnetically labeled agarose beads.**

(**A**) Time-dependent spatial distributions of magnetically labeled agarose beads before and after shrinking. Beads were placed in wells containing a central neodymium magnet, and the distance of each bead from the magnet center was quantified over time. The shaded region indicates the magnet area. Beads before shrinking gradually accumulated near the magnet over 50 min, whereas shrunken beads rapidly shifted toward the magnet and became concentrated near the center within 15 min. These results show that bead shrinking improves magnetic attraction kinetics.

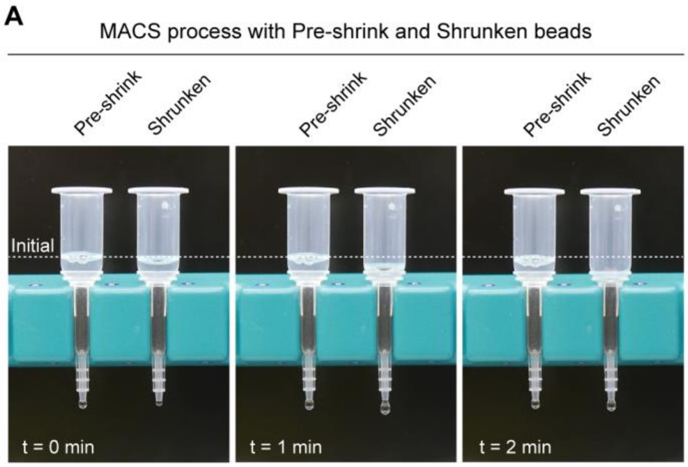


**Figure S4. Bead shrinking prevents clogging during MACS column processing.**

(**A**) Representative photographs of commercial MACS columns loaded with agarose beads before and after shrinking. Beads before shrinking rapidly accumulated at the column inlet and clogged the column, preventing efficient flow-through and subsequent washing. In contrast, shrunken beads passed through the column without observable clogging during the MACS procedure, demonstrating that bead size reduction is required for compatibility with commercial magnetic columns.

A

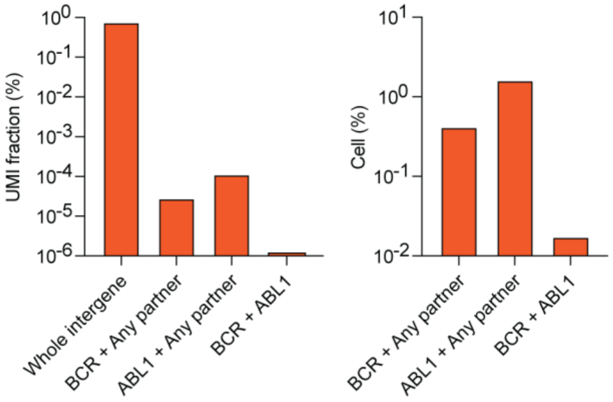


**Figure S5. Prevalence of putative intergenic chimeric transcripts in the publicly available K562 single-cell RNA-sequencing dataset.**

(**A**) Fractions of unique cell barcode–UMI pairs associated with whole-intergene, BCR–any partner, ABL1–any partner, and BCR::ABL1 chimeric alignments (left), and percentages of the 5,768 called cells containing BCR–any partner, ABL1–any partner, or BCR–ABL1 signals (right).

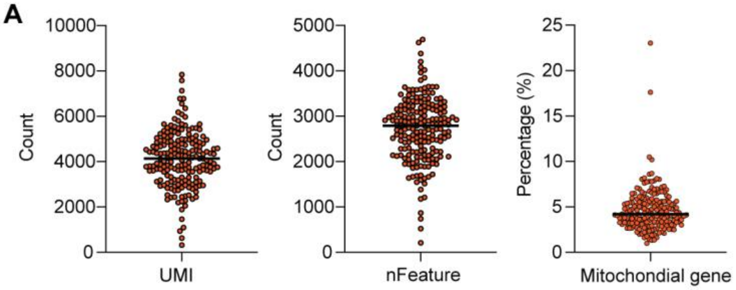


**Figure S6**. **Quality-control metrics of magniFIND-seq-isolated K562-r single-cell transcriptomes.**

(**A**) Distributions of total UMI counts, numbers of detected genes (nFeature), and percentages of mitochondrial transcripts across the 182 magniFIND-seq-isolated cells that passed quality-control filtering. Each point represents an individual cell, and horizontal lines indicate median values. Most cells contained several thousand UMIs and detected genes while maintaining low mitochondrial transcript fractions.
